# Automating Biomedical Knowledge Graph Construction For Context-Aware Scientific Inference

**DOI:** 10.64898/2026.01.14.699420

**Authors:** Yikai Zheng, Wanquan Liu, Bi Zeng, Yichun Feng, Xiawei Du, Lu Zhou, Yixue Li

**Affiliations:** School of Intelligent Systems Engineering, Sun Yat-sen University, No. 66, Gongchang Road, Guangming District, Shenzhen, 518107, Guangdong, China.; Guangzhou National Laboratory, No. 9 XingDaoHuanBei Road, Guangzhou International Bio Island, Guangzhou, 510005, China.; School of Computer Science and Technology, Guangdong University of Technology, Guangzhou, 510000, China.; School of Advanced Interdisciplinary Sciences, University of Chinese Academy of Sciences, 380, Huaibei Town, Huairou District, Beijing, 100049, Beijing, China.; Department of Computer Science and Technology, Tsinghua University, Haidian District, Beijing, 100084, P. R. China.

**Keywords:** Biomedical Knowledge Graph, Composite triple, Context-aware Inference, Large Language Models, Open Information Extraction, Self-training

## Abstract

Biomedical interactions are inherently dynamic, often shifting or even reversing under specific physiological states. However, existing extraction methods simplify these complex mechanisms into context-agnostic binary associations, resulting in semantic loss and contradictory evidence. Here, we present AutoBioKG, an end-to-end framework that constructs context-aware knowledge graphs by leveraging composite triples to encode environmental conditions and entity attributes alongside core relationships. Powered by an open information extraction model trained on BioOpenIE and further refined through self-training with pseudo-labels from unlabeled literature, the framework exhibits broad generalization. Notably, AutoBioKG achieved the highest zero-shot F1 across DDI, ChemProt, and BioRED, outperforming the best-performing baseline on each benchmark by 3.6–17.8 percentage points. Furthermore, AutoBioKG-derived graphs outperformed existing approaches on yes/no, factoid, and list questions in the BioASQ biomedical question-answering evaluation, particularly for queries requiring fine-grained contextual information. Together, these results support AutoBioKG as a scalable framework for transforming unstructured literature into structured, context-aware biomedical knowledge.

## 1 Introduction

Biomedical knowledge, embedded within millions of research articles, holds transformative potential for accelerating drug discovery, advancing precision medicine and elucidating disease mechanisms. However, this valuable knowledge predominantly exists in unstructured textual form, limiting direct utilization and systematic analysis. Knowledge graphs (KGs), which represent entities and their complex relationships in a structured format, have emerged as a powerful solution for organizing and leveraging biomedical knowledge [1–4]. By mapping drug-target-disease interaction networks [1] and supporting personalized medicine [2, 3], these structured representations have become indispensable infrastructure for modern computational biology [4, 5].

Despite these advancements, a fundamental disconnect persists between the richness of biological literature and the expressiveness of current extraction methods. Traditional Information Extraction (IE) methods rely on predefined schemas, facing severe scalability challenges when processing the rapidly emerging relationships in biomedical literature [6–8]. While Open Information Extraction (OIE) offers a schema-free alternative [9–13], the prevailing subject-predicate-object triples employed by these systems reduce complex biological mechanisms into static, context-agnostic snapshots. For instance, biological interactions are inherently dynamic and context-dependent, as a relationship may exist only under specific cellular states. Simplifying the statement “Under hypoxic conditions, PTPMeg2 dephosphorylates its down- stream target STAT3 at the Tyr705 site” into the triple (PTPMeg2, dephosphorylates, STAT3) removes the hypoxic condition and the Tyr705 attribute, obscuring the context in which the interaction was reported. Crucially, this is not an isolated edge case. Our quantitative analysis reveals that over 72% of biomedical documents contain such conditional or attribute-dependent relationships, suggesting that conventional triple representations systematically discard the contextual constraints essential for precision medicine.

However, the development of models capable of extracting such complex, structured knowledge faces a fundamental bottleneck of data availability. Existing biomedical relation extraction datasets, such as BioRED [14] and DDI [15], are mainly designed for tasks with predefined relation types. They lack systematic annotations for contextual conditions and entity attributes, making it extremely challenging to train extraction models that can handle open-domain scenarios and capture fine-grained semantics. Furthermore, the extracted triples cannot be used directly for knowledge graph construction. Inconsistent entity naming and diverse relational expressions cause the raw output to contain significant redundancy and semantic ambiguity [16, 17]. Moreover, the lack of systematic normalization and alignment mechanisms makes it difficult to integrate these triples into a coherent knowledge graph. These challenges motivate an automated pipeline that links information extraction with knowledge fusion and graph construction. Recently, the integration of large language models (LLMs) and Knowledge Graphs has garnered significant attention as a promising roadmap to address these scalability and reasoning challenges [18–20]. Although recent frameworks for automated knowledge graph construction exist [21–23], they often struggle to capture complex contextual dependencies or lack effective mechanisms for knowledge fusion.

To address these challenges, we present AutoBioKG, an end-to-end framework for automatically constructing context-aware open-domain biomedical knowledge graphs from unstructured text. Central to this framework is a novel representation called composite triples, which structurally encodes contextual conditions and entity attributes to capture the context-dependent nature of biological mechanisms. Unlike static representations, this schema renders the knowledge graph as a context-aware topology where relationships adapt based on entity attributes and environmental conditions. To enable this representation, we developed a specialized open information extraction model trained on BioOpenIE, a high-quality dataset we constructed through multilayered automatic annotation strategies. Beyond BioOpenIE, we extend the training data through self-training on unlabeled biomedical literature. The same documents are processed once by the multitask seed model and again by separately trained single-task models, producing two sets of composite triples. Their outputs are merged before final training, increasing the range of biomedical relations represented in the pseudo-labeled data. Following extraction, AutoBioKG integrates a knowledge fusion module that ensures semantic consistency and standardization through entity normalization and dynamic alignment of novel relationships. The resulting graph is automatically instantiated in a Neo4j database, supporting complex queries.

Applying this framework, we successfully constructed high-quality knowledge graphs from biomedical literature and evaluated the framework across relation extraction and biomedical question answering tasks. Our self-trained OIE model achieved the highest F1 among the evaluated LLMs in zero-shot testing on DDI, ChemProt, and BioRED without task-specific fine-tuning, outperforming the best-performing baseline on each benchmark by 3.6, 17.8, and 6.2 percentage points, respectively. On the BioASQ benchmark, knowledge graphs constructed by AutoBioKG achieved 34.1% F-measure on list questions and 82.7% macro F1 on yes/no questions, out- performing the compared automated construction methods on both question types. These results demonstrate the potential of AutoBioKG to transform biomedical literature into context-rich structured knowledge through an automated pipeline, providing useful evidence for downstream biomedical question answering.

## 2 Methodology

### 2.1 The overview of AutoBioKG

AutoBioKG aims to automatically construct high-quality open-domain knowledge graphs from unstructured biomedical literature through end-to-end information extraction and knowledge fusion (Fig. 1a). The framework combines a self-trained OIE model with an automated knowledge fusion module.

**Fig. 1:**
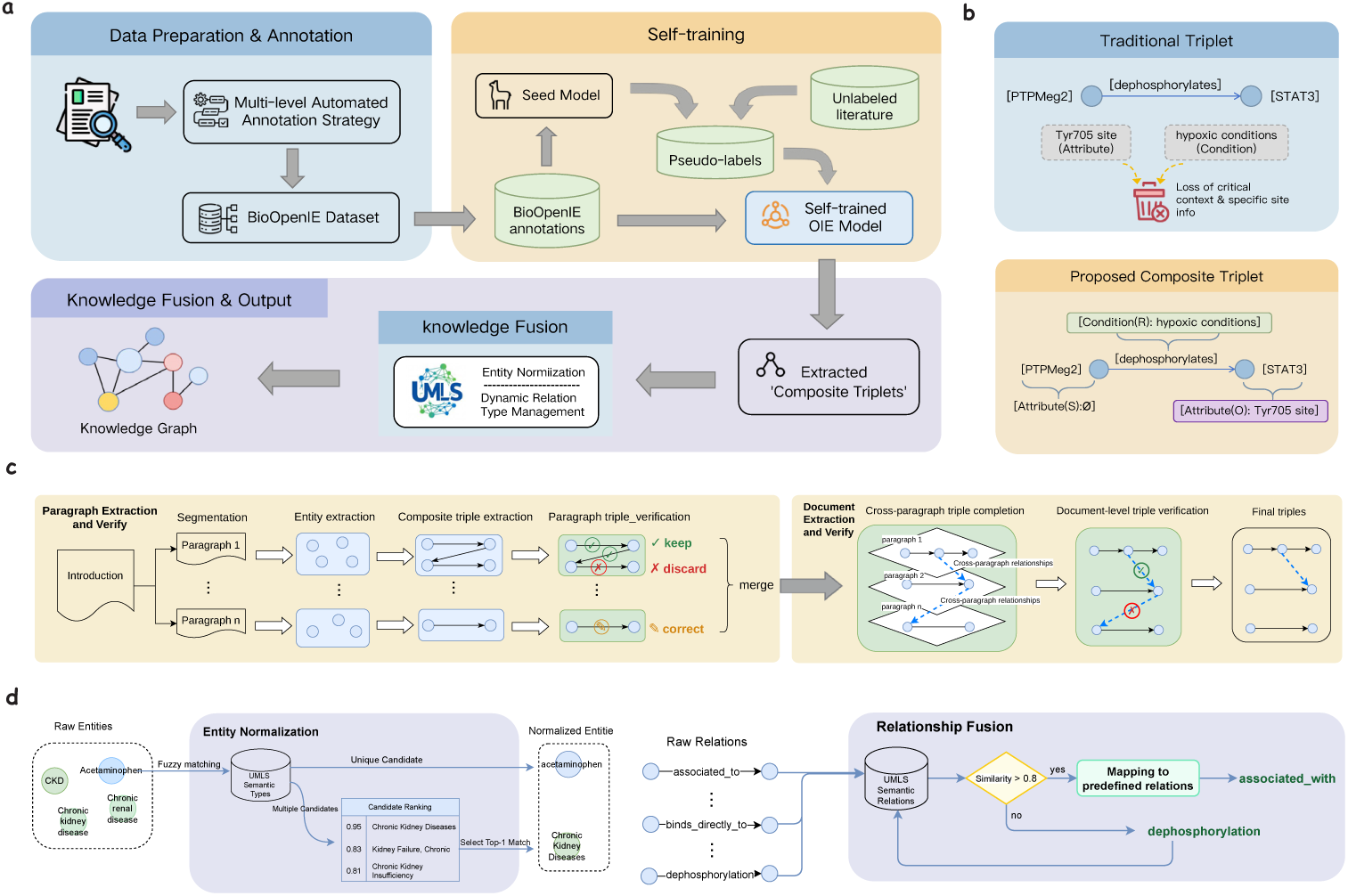
Overview of the AutoBioKG framework and construction methodology. **a,** Schematic overview of the automated knowledge graph construction pipeline. Unstructured biomedical literature is processed through an automated extraction and knowledge-fusion workflow to construct the graph. **b,** Comparison of knowledge representation schemas. Unlike conventional subject-predicate-object triples, composite triples explicitly represent entity attributes and relation conditions. **c,** Workflow for the LLM-powered construction of the BioOpenIE dataset. The annotation process progresses from semantic segmentation and entity extraction to cross-paragraph triple completion and document-level verification. **d,** Knowledge fusion and semantic standardization module. Raw extracted entities are normalized to UMLS concepts through exact candidate retrieval and SapBERT-based semantic disambiguation, while relations are dynamically aligned to unified schemas based on embedding similarity.

At the core of AutoBioKG lies a knowledge representation termed composite triples, which extends the conventional subject-predicate-object structure by explicitly encoding entity-specific attributes and contextual conditions, as shown in Fig. 1b. This representation captures semantic information that conventional triples do not explicitly retain, including tissue context in gene-disease associations and experimental conditions in drug interactions. To operationalize this representation, we constructed BioOpenIE, an open information extraction dataset for biomedical literature. Built upon 500 biomedical research papers through a multi-stage automated annotation pipeline (Fig. 1c), the dataset comprises 5,129 composite triples. Notably, 24.37% of these triples contain enriched attributes or conditions, while such complex dependencies permeate 72.25% of the analyzed documents, highlighting that contextual information provides crucial training signals for learning complex biomedical mechanisms rather than being a marginal feature.

Starting from the BioOpenIE-trained seed model, self-training incorporates pseudo-labeled examples from unlabeled biomedical literature. The same documents are processed under two model configurations, and the combined outputs are used to train a single resulting OIE model for downstream extraction. Following extraction, the knowledge fusion module standardizes entities and relations through UMLS-based canonicalization and dynamic relation alignment (Fig. 1d), ensuring semantic consistency while accommodating emerging relations in the literature. The resulting knowledge is automatically stored in a Neo4j graph database, forming a structured and queryable resource.

### 2.2 Open information extraction via composite triples

The proposed AutoBioKG framework employs a specialized open information extraction model designed to capture the dynamic nature of complex biomedical semantics via composite triples. To extend training beyond BioOpenIE, we apply self-training to unlabeled biomedical text. The self-trained OIE model is obtained in two stages. We first train a seed model on BioOpenIE using supervised fine-tuning. Next, two model configurations process the same unlabeled documents, and their merged triple sets are used to retrain the seed model. This section details the composite triple schema, model architecture, and training procedure.

### Composite Triple Representation

Conventional open information extraction systems represent knowledge as subject-predicate-object (SPO) triples. To overcome the semantic loss discussed in the introduction, our framework adopts a composite triple schema designed to capture dynamic biological mechanisms. Using the running example “Under hypoxic conditions, PTPMeg2 dephosphorylates STAT3 at Tyr705 site”, where conventional OIE would extract the triple (PTPMeg2, dephosphorylates, STAT3), our schema preserves the structural integrity by explicitly mapping *context* and *state* into dedicated slots. Specifically, “hypoxic conditions” is encoded as the *Condition* variable determining the relation’s validity, while “Tyr705” is captured as an *Attribute* defining the object’s state. Formally, each composite triple is defined as:

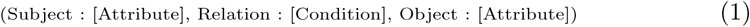

where the subject and object represent core entities and the relation denotes their semantic interaction. Attribute and Condition are optional slots. Attribute captures entity-specific attributes such as gene mutations (EGFR [L858R]) and protein sites (STAT3 [Tyr705]). Condition encodes the contextual circumstances under which the relation holds, such as experimental settings (in vitro), tissue contexts (in lung tissue) or temporal constraints (during G1 phase). Our schema binds these elements to define the relationship as a dynamic function of the interacting entity states and environmental contexts. This formalism enables the representation of a state-dependent topology, providing a rigorous foundation for reasoning about dynamic biological mechanisms.

### Model Architecture and Training Objective

Our open information extraction model is built upon LLaMA3.1-8B. This open-source large language model features 8 billion parameters and was selected for its robust instruction-following capability and efficiency in processing large-scale literature. The model adopts a standard decoder-only Transformer architecture comprising 32 layers, 32 heads, a hidden dimension of 4,096, and an intermediate dimension of 14,336. Given an input sentence *x* = *{x*_1_*, x*_2_*, …, x_n_}*, the model autoregressively generates a composite triple sequence *y* = *{y*_1_*, y*_2_*, …, y_m_}* by maximizing conditional probability:

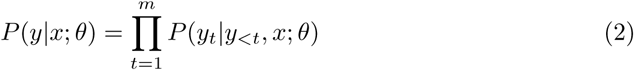

where *θ* represents model parameters and *y_<t_* denotes previously generated tokens. Instead of full-parameter tuning, we employ Low-Rank Adaptation [24] (LoRA), where *θ* consists only of the low-rank decomposition matrices injected into the attention layers, while the original pre-trained weights remain frozen. During supervised fine-tuning, the model is optimized to generate composite triples by minimizing negative log-likelihood:

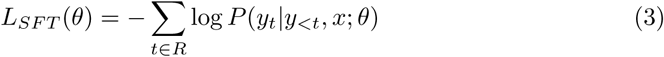

where *x* represents input instructions and documents, *y* = *{y*_1_*, y*_2_*, …, y_m_}* represents the target response sequence, and *R* denotes the set of response token positions. Instruction tokens are masked in the loss computation to ensure the model focuses on learning response generation.

### Self-training

To extend the BioOpenIE training set, we generate additional examples from unlabeled biomedical text under two model configurations. Using two configurations reduces dependence on the extraction patterns of a single generator and broadens the relations represented in the training data.

For each unlabeled document, both configurations perform composite triple extraction, cross-paragraph triple completion, and multi-layer verification. In the first configuration, the multitask seed model performs all three stages. In the second, extraction, completion, and verification are handled by separately trained single-task models.

Within each document, the two outputs are deduplicated and merged at the triple level. All retained pseudo-labels receive equal weight and are combined with the original BioOpenIE annotations. Under the same fine-tuning settings, the multitask seed model is then retrained. Single-task models are used only to generate training data; downstream extraction relies on the resulting OIE model.

### Training Pipeline

The seed model was trained on BioOpenIE using the AdamW optimizer for supervised fine-tuning with a learning rate of 1 *×* 10*^−^*^4^ and weight decay of 0.01. We configured the LoRA adapter with a rank *r* = 64, a scaling factor *α* = 128, and a dropout rate of 0.05. Training utilized one NVIDIA A100 GPU with a batch size of 16 and adopted half-precision FP16 computation. The model was trained for 3 epochs. To implement self-training, we sampled 300 introductions from biomedical PubMed articles and processed each document under both model configurations. These unlabeled articles were disjoint from the 500 articles used to construct BioOpenIE and from the documents included in the downstream relation-extraction evaluation sets. After document-level deduplication and triple-level merging, the resulting pseudo-labeled examples were added to the BioOpenIE training data. The seed model was then fine-tuned for another three epochs using the same optimizer, LoRA configuration, and batch settings. This model was used in all subsequent extraction experiments.

### 2.3 The BioOpenIE dataset construction

To provide supervision for context-aware open information extraction, we constructed BioOpenIE, a biomedical open information extraction dataset based on the composite triple representation. The dataset contains 5,129 composite triples generated from 500 biomedical research articles through a multi-stage annotation pipeline using multiple large language models. This design provides training examples in which relation extraction is learned together with the attributes and conditions needed to preserve biomedical context.

Literature Corpus Selection and Preprocessing. We curated 500 peer-reviewed research articles from the PubMed database, selected to ensure comprehensive coverage of diverse biomedical domains. MinerU [25] was used to convert the articles into structured Markdown while preserving document hierarchy. Recognizing that the Introduction section contains the highest density of established biological relations, we programmatically extracted this section from each article by matching common header patterns. This process yielded an average of 497 words per article for subsequent annotation.

Multi-stage Automated Annotation Pipeline. To generate high-quality annotations without extensive human intervention, we designed a six-step pipeline orchestrating multiple large language models to execute complementary tasks as shown in Fig. 1c. The first four steps operate at the paragraph level, after which the paragraph-level outputs are merged for cross-paragraph triple completion and document-level triple verification. Detailed prompt templates for all six steps are provided in Appendix C. Step 1: Semantic Segmentation and Coreference Resolution. The first step utilizes GPT-4.1 for document preprocessing. For the Introduction section of each article, the model partitions the text into 3 to 5 semantically coherent paragraphs, ensuring that each paragraph contains an independent conceptual unit suitable for relation extraction. The same step resolves coreferences by replacing pronominal references such as “this protein” with the corresponding entity names, reducing ambiguity in subsequent extraction.

Step 2: Entity Extraction. Guided by a taxonomy of 23 UMLS [26] semantic types relevant to biomedical research as detailed in Table 5, GPT-4.1 identifies biomedical entities within each paragraph. The prompt enforces explicit constraints requiring the extraction of specific terms rather than generic categories. For example, it requires extracting “lactase” instead of “enzyme”. It also requires abbreviated entities to be represented together with their full names and excludes overly general descriptors. This constrained extraction strategy ensures entities are sufficiently fine-grained for precise relation modeling.

Step 3: Composite Triple Extraction. Using the tokenized text and identified entities as input, GPT-4.1 extracts composite triples following the schema defined in Equation (1). The prompt explicitly specifies the composite structure and provides annotated examples demonstrating the identification of entity attributes such as gene mutations and drug dosages alongside relational conditions such as experimental backgrounds and cellular environments. Crucially, the prompt emphasizes extracting biologically significant interactions while excluding attributive relations like “is” or “characterized by”, experimental setting descriptions like “measured by”, and uncertain statements like “may result in”.

Steps 4–6: Multi-layer Verification and Completion. DeepSeek-R1 [27] executes three layers of quality control. First, paragraph-level triple verification ensures that each extracted triple is grounded in the source paragraph by checking whether its entities and relation are supported by the text. Second, cross-paragraph triple completion identifies interactions in which the subject and object appear in different sentences and may therefore be missed during paragraph-level extraction. This step recovers relations that depend on evidence distributed across sentence boundaries. Third, document-level triple verification reviews the combined triple set against the full Introduction to identify unsupported relations, contradictions, and redundancies. Triples are corrected when the source text supports a revision and discarded when no supported correction can be made.

Data Splitting and Quality Validation. The dataset was partitioned based on source literature into a training set comprising 300 articles and a test set comprising 200 articles. To estimate annotation quality, we randomly sampled 100 documents from the test set. A PhD-level computational biologist and Gemini 3 Pro [28] independently assessed the extracted triples as correct or incorrect according to the same evaluation criteria. The two evaluators reached a raw agreement of 94.2%. Disagreements were subsequently reviewed to establish a consensus reference, against which the annotation pipeline achieved a triple-level accuracy of 87.1%.

### 2.4 Knowledge fusion and graph construction

Following open information extraction, we implement a multi-stage knowledge fusion pipeline to ensure the semantic consistency, normalization, and structural integrity of the final knowledge graph. The pipeline integrates entity normalization via Unified Medical Language System (UMLS) concept linking, relation alignment via dynamic vocabulary expansion, and automated storage in a Neo4j graph database.

Entity Normalization. To standardize entities extracted from diverse literature sources, we associate each entity mention with canonical concepts in the UMLS, a comprehensive metathesaurus integrating over 200 biomedical vocabularies. For each extracted entity *e*, we query the UMLS API to retrieve candidate Concept Unique Identifiers (CUI). When a single CUI is returned, we establish a direct mapping. For cases yielding multiple candidate CUIs, we employ the SapBERT [29] model for semantic disambiguation to calculate contextual similarity. Specifically, we utilize the entity as input and compute the cosine similarity based on the canonical name and definition of each candidate CUI. If the similarity score exceeds a preset threshold *θ* of 0.8, we map the entity to the candidate with the highest similarity score. Entities failing to reach this threshold retain their original text form as novel entities not yet incorporated into UMLS, preserving terminology emerging in recent literature.

Relation Alignment and Dynamic Vocabulary Expansion. To unify the diverse relation expressions generated by open information extraction, we maintain a dynamic standard relation vocabulary initialized with predefined relation types from the UMLS Semantic Network. For each extracted relation *r*, we employ the BGE-m3 [30] encoder to generate its embedding and compute its cosine similarity with all standard relations in the current vocabulary as defined in:

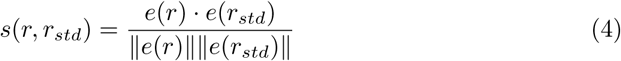

where *e*(*r*) and *e*(*r_std_*) represent the embeddings of the extracted and standard relations, respectively. If the maximum similarity score exceeds the relation threshold *θ_relation_* = 0.8, *r* is mapped to the best-matching standard relation. Otherwise, the relation is standardized and added to the relation vocabulary as a new type. This match-or-add strategy ensures consistency in relation expression. Furthermore, it enables the knowledge graph to continuously learn and expand its knowledge system by automatically identifying and accommodating novel relations emerging in the literature.

## 3 Experiments

### 3.1 Datasets

To comprehensively assess both extraction quality and practical utility, we evaluated AutoBioKG across multiple benchmarks encompassing open-domain information extraction, closed-domain relation extraction, and biomedical question answering.

Open-domain Information Extraction. The core extraction model was trained and evaluated on BioOpenIE. The dataset contains 5,129 composite triples generated from 500 peer-reviewed biomedical research articles. To prevent relations from the same source article appearing in both partitions, the split was performed at the article level, with 300 articles used for training and 200 for testing.

Closed-domain Relation Extraction. To assess the model’s zero-shot transfer capability to unseen schemas, we employed three established biomedical benchmarks. The DDI [15] dataset comprises 1,025 documents from DrugBank and MEDLINE annotated for drug-drug interactions, with one entity type and four relation types. The ChemProt [31] dataset, annotated by domain experts, comprises 1,820 PubMed abstracts annotated for chemical-protein interactions, with two entity types and 23 relation types. The BioRED [14] dataset serves as a larger-scale corpus for extracting relationships between biomedical entities such as genes and diseases from PubMed abstracts, covering a broader range of relation types.

Biomedical question answering. We employed the BioASQ Task 12b (2024) benchmark to evaluate the practical utility of knowledge graphs constructed by AutoBioKG. The original dataset contains 5,046 questions paired with gold-standard answers. However, the prevalence of clinical guideline queries and epidemiological statistical questions limits the suitability of the raw data for assessing the graph’s mechanistic reasoning capabilities. To address this, we implemented an LLM-based filter to retain questions involving biomedical interactions and remove purely definitional or procedural queries. The resulting evaluation set contained 784 questions: 200 yes/no, 200 factoid, 184 list, and 200 summary questions. For each question, the associated expert snippets were concatenated into a question-specific evidence package, which served as the source text for graph construction.

### 3.2 Evaluation metrics

The performance of the open-domain information extraction task is quantified by precision, recall, and F1 values. Given the inherent linguistic expression diversity in open-domain environments, traditional methods often erroneously penalize semantically equivalent expressions, thereby underestimating model performance. To address this limitation, we employed a semantic similarity matching framework. Under this framework, if a predicted triple is semantically consistent with a true triple, it is regarded as a true positive. Specifically, each composite triple is converted into a coherent natural language sentence and encoded into high-dimensional vectors utilizing a pre-trained BERT [32] model. Then, by calculating the cosine similarity of the respective vectors of the predicted triple and true triple, the semantic relevance between them is quantified. If the similarity score exceeds a preset threshold *θ* = 0.85, the match is successful. Based on this matching scheme, precision is defined as the proportion of successfully matched predicted triples, while recall is defined as the proportion of successfully matched true triples. The F1 score is the harmonic mean of precision and recall, providing a comprehensive measure of model performance.

The evaluation scheme for the biomedical question answering task adopts BioASQ evaluation metrics corresponding to the four question types. For “Yes/No” questions, accuracy and macro F1 are used to evaluate binary classification performance, with macro F1, defined as the unweighted average of the class-wise F1 scores, serving as the primary evaluation metric. For “Factoid” questions, Strict Accuracy counts a question as correct when a gold-standard entity or its synonym appears in the first returned position, whereas Lenient Accuracy counts it as correct when a gold-standard entity or its synonym appears anywhere in the returned candidate list. The performance of “List” questions is evaluated using mean precision, recall, and F-measure, which quantify the quality of the predicted list. Finally, the generated “Summary” answers are evaluated using the ROUGE [33] metric system, including ROUGE-2 recall and F1, as well as ROUGE-SU4 recall and F1, to assess overlap with the reference summaries.

### 3.3 Self-training improves extraction performance

To assess the efficacy of self-training, we benchmarked the self-trained OIE model against the initial seed model. Our evaluation encompassed three widely recognized public datasets, namely DDI [15], ChemProt [31], and BioRED [14], alongside our BioOpenIE dataset. Both models were evaluated using a semantic similarity matching framework that assesses extraction quality through BERT-encoded semantic similarity rather than exact string matching.

Across all four datasets, the self-trained model consistently outperformed the seed model, as shown in Fig. 2b. Specifically, F1 improvements ranged from 4.2 to 7.6 percentage points. The largest gains were observed on BioRED, ChemProt, and BioOpenIE, where F1 increased by 7.6, 7.4, and 6.8 percentage points, respectively. Notably, while BioOpenIE achieved the highest absolute score, as expected for the in-domain evaluation set, the gains on ChemProt and BioRED were comparable to or greater than the gain observed on BioOpenIE. These results show that the benefit of self-training extended beyond the BioOpenIE test set to public benchmarks with different relation schemas.

**Fig. 2:**
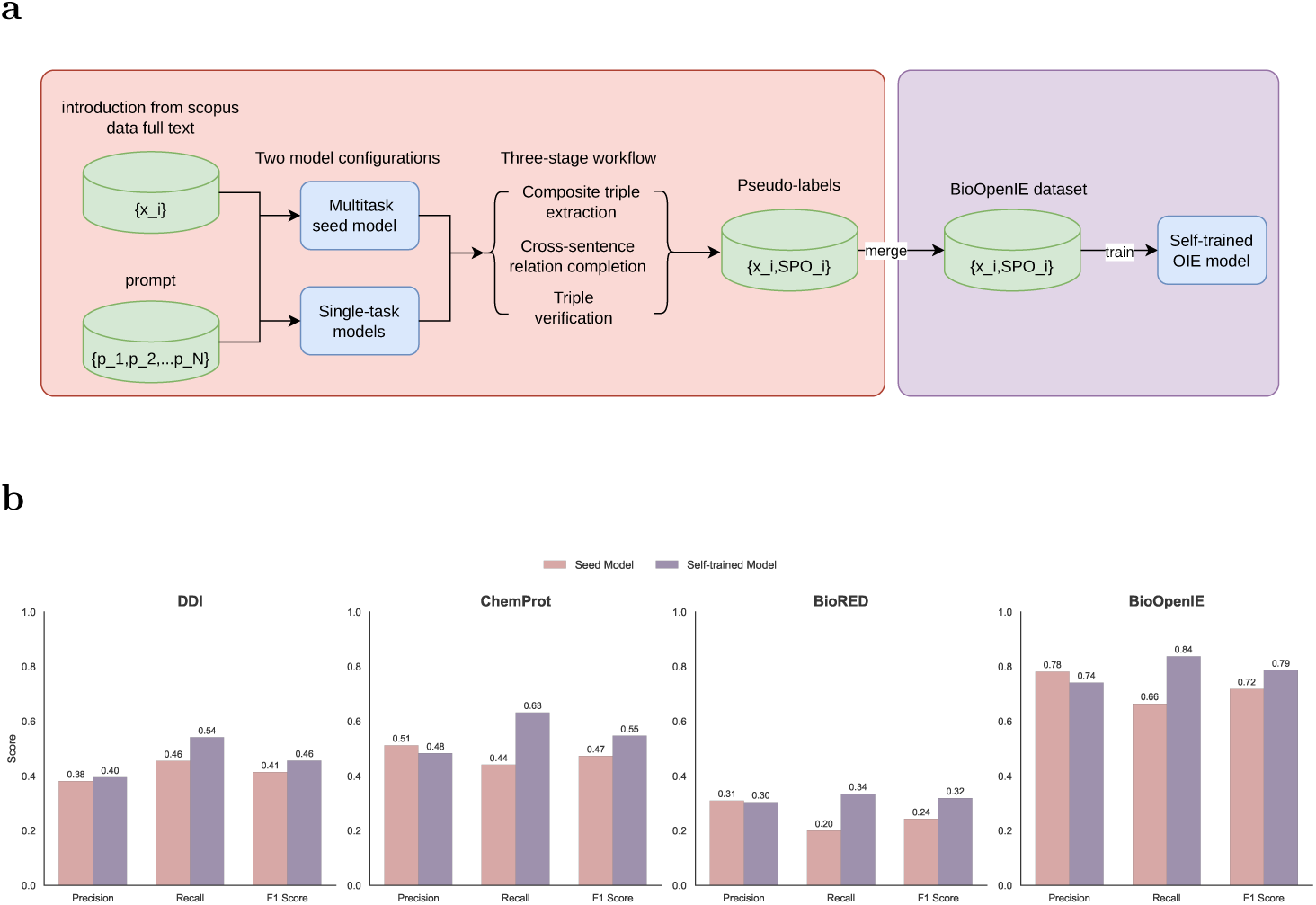
Self-training workflow and extraction performance. **a,** Self-training workflow using two model configurations on unlabeled biomedical literature. The multitask seed model and separately trained single-task models generate composite triples through extraction, cross-paragraph triple completion, and triple verification. Their outputs are merged into pseudo-labels and combined with BioOpenIE to train the self-trained OIE model. **b,** Precision, recall, and F1 of the seed and self-trained OIE models on DDI, ChemProt, BioRED, and BioOpenIE.

### 3.4 The self-trained OIE model demonstrates strong generalization to unseen relation schemas

Having validated the efficacy of self-training, we next assessed whether the self-trained OIE model could generalize to predefined relation extraction tasks in a zero-shot setting. We evaluated the model on DDI [15], ChemProt [31], and BioRED [14] without task-specific fine-tuning on any of these datasets. This evaluation is critical for assessing the practical utility of the model, as biomedical applications often require the extraction of relations conforming to unseen schemas. We benchmarked our model against six LLM baseline configurations, including GPT-4o [34], GPT-4.1 [35], GPT-5 mini [36], LLaMA3.1-8B [37], Qwen3-8B [38] in non-thinking mode, and Qwen3-8B in thinking mode.

Our model achieved the highest F1 across all three predefined relation extraction benchmarks, as shown in Table 1. The advantage was most pronounced on ChemProt, where the margin over the strongest baseline reached 17.8 percentage points. This F1 advantage was accompanied by the highest precision across all three benchmarks and the highest recall on ChemProt and BioRED.

**Table 1:** Document-level relation triplet extraction performance on the public evaluation datasets.

| Model | DDI |  |  | ChemProt |  |  | BioRED |  |  |
| --- | --- | --- | --- | --- | --- | --- | --- | --- | --- |
|  | Precision | Recall | F1 | Precision | Recall | F1 | Precision | Recall | F1 |
| LLaMA3.1-8B | 0.294 | **0.619** | 0.398 | 0.291 | 0.505 | 0.369 | 0.181 | 0.304 | 0.227 |
| Qwen3-8B (no thinking) | 0.283 | 0.529 | 0.369 | 0.291 | 0.451 | 0.354 | 0.216 | 0.318 | 0.257 |
| Qwen3-8B (thinking) | 0.360 | 0.503 | 0.420 | 0.323 | 0.370 | 0.345 | 0.256 | 0.248 | 0.252 |
| GPT-4o | 0.334 | 0.560 | 0.418 | 0.315 | 0.433 | 0.364 | 0.166 | 0.263 | 0.204 |
| GPT-4.1 | 0.298 | 0.585 | 0.395 | 0.287 | 0.489 | 0.362 | 0.142 | 0.230 | 0.176 |
| GPT-5 mini | 0.234 | 0.520 | 0.323 | 0.229 | 0.400 | 0.292 | 0.116 | 0.198 | 0.146 |
| **Our model** | **0.395** | 0.541 | **0.456** | **0.483** | **0.632** | **0.547** | **0.304** | **0.335** | **0.319** |

These results collectively demonstrate that extraction capabilities developed through open-domain training on composite triples can successfully transfer to novel predefined relation schemas. This generalization may reflect the alignment between the composite triple representation and the semantic knowledge encoded in the pretrained language model, enabling the model to capture biomedical relations beyond the characteristics of individual datasets.

### 3.5 Composite triples preserve contextual distinctions in biomedical relations

Biomedical relations that share the same core entities can differ in treatment or patient context. Figure 3a illustrates this distinction in two treatment-related examples. In the tamoxifen example, the graph retains the condition in women treated with tamoxifen together with the CYP2D6-mediated pathway from tamoxifen to endoxifen. In the olaparib example, germline BRCA1/2 mutation and platinum resistance distinguish patient contexts associated with the same drug-disease relation. In both cases, the

**Fig. 3:**
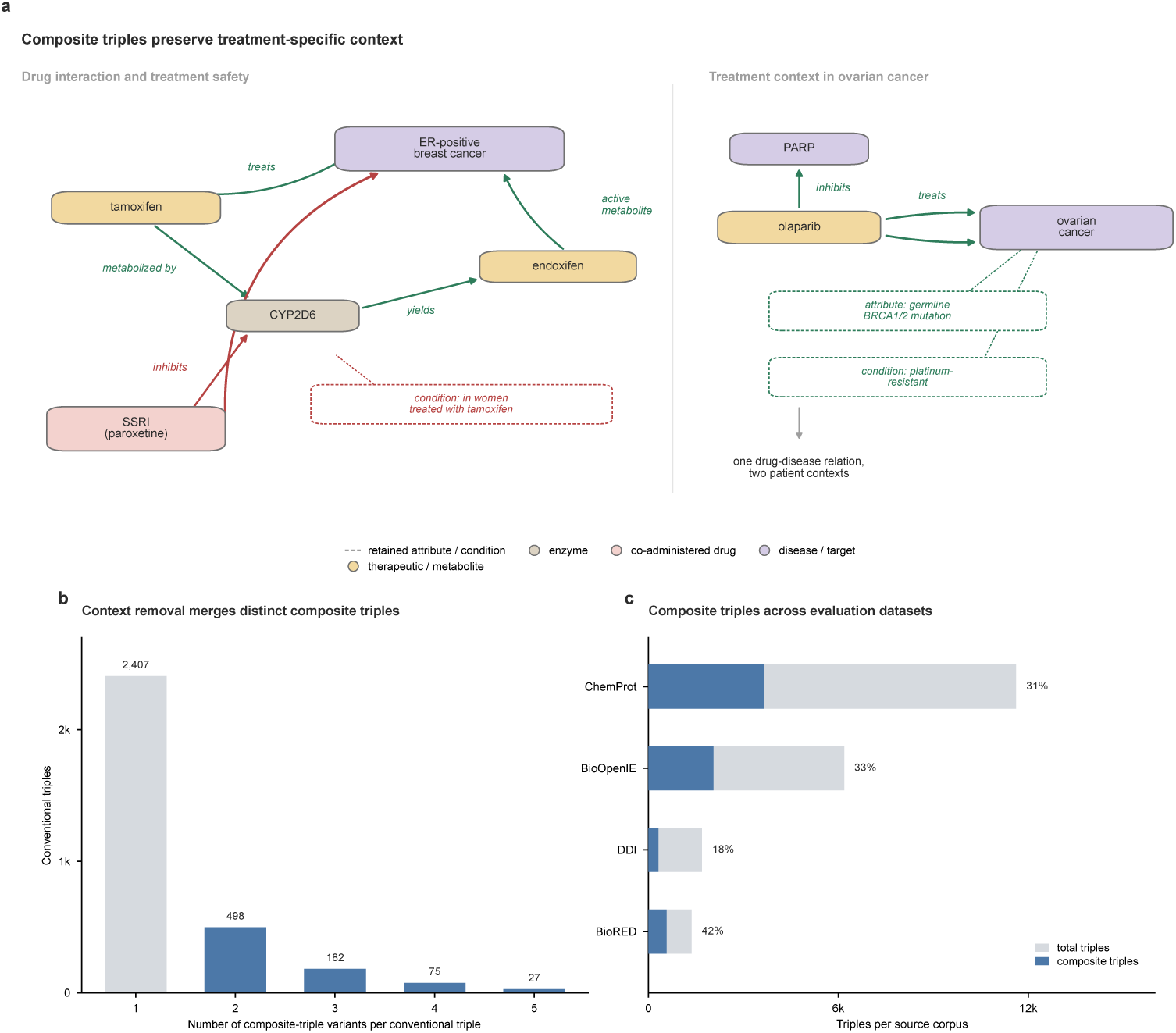
Composite triples preserve contextual distinctions in biomedical relations. **a,** Treatment-related examples showing how entity attributes and relation conditions retain treatment and patient context that is omitted from conventional triples. **b,** Distribution of composite triple variants that map to the same conventional triple after Attribute and Condition fields are removed. Of 3,189 conventional triples, 782 correspond to two or more distinct composite triple variants. **c,** Total extracted triples and composite triples across ChemProt, BioOpenIE, DDI, and BioRED. Percentages denote the proportion of extracted triples containing at least one Attribute or Condition field.

Attribute and Condition fields preserve information that would be omitted from a conventional subject-predicate-object representation.

Removing this contextual information caused distinct composite triples to collapse into the same conventional triple. Among the 3,189 conventional triples summarized in Fig. 3b, 782 corresponded to two or more distinct composite triple variants. Most of these cases involved two variants, although some conventional triples merged as many as five. This many-to-one mapping shows directly that removing Attribute and Condition fields can erase distinctions that are retained by the composite triple representation.

The same representation was observed across all four evaluation corpora. Composite triples accounted for 31% of extracted triples in ChemProt, 33% in BioOpenIE, 18% in DDI, and 42% in BioRED as shown in Fig. 3c. Although their prevalence varied across relation domains, composite triples were recovered in each external evaluation dataset, indicating that explicit attributes and conditions were not confined to BioOpenIE.

### 3.6 AutoBioKG-constructed knowledge graph enhances biomedical question answering

To evaluate the practical utility of the AutoBioKG knowledge graph, we conducted tests using BioASQ [39] as a benchmark. BioASQ is a biomedical question answering benchmark containing four question types, namely, Yes/No, Factoid, List, and Summary. We integrated AutoBioKG and three baseline methods, specifically AutoKG [21], EDC [23], and iText2KG [22], into a common Knowledge Graph Retrieval-Augmented Generation (KG-RAG [40]) pipeline. In this setting, a Large Language Model (LLM) identifies key entities in the query, retrieves relevant facts from the knowledge graph, and generates the final answer. Detailed implementation specifications, including graph-specific entity-matching settings, hyperparameter settings, and prompt templates, are provided in Appendix B. The methods share the same answer-generation and context-pruning procedure, reducing downstream implementation differences and focusing the comparison on the information represented in the resulting graphs.

#### 3.6.1 Quantitative performance analysis across BioASQ tasks

AutoBioKG achieved the strongest overall performance on yes/no, factoid, and list questions, while remaining competitive on summary questions, as shown in Fig. 4a. The largest advantage was observed for list questions, while substantial gains were also seen on yes/no questions. For yes/no questions, AutoBioKG reached an accuracy of 87.0% and a macro F1 score of 82.7%, outperforming the compared graph-construction methods. This performance is consistent with the semantic standardization introduced by the knowledge fusion module. Entity normalization reduces redundancy across synonymous biomedical concepts, while relation alignment promotes consistent relation expressions in the resulting graph. Such standardization can reduce ambiguity in the evidence retrieved for binary questions.

**Fig. 4:**
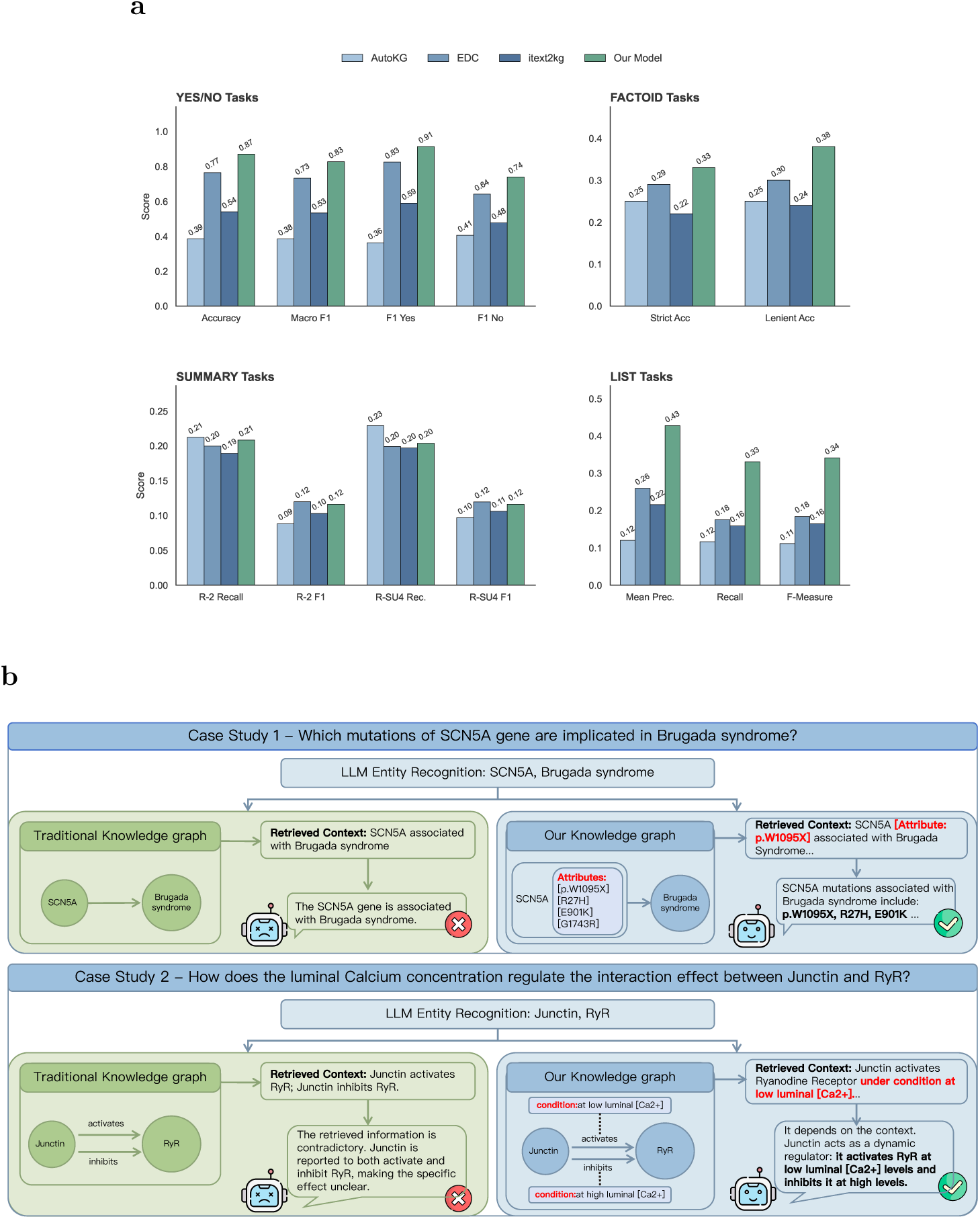
Quantitative and qualitative evaluation of knowledge graph-enhanced question answering. **a,** Performance comparison on the BioASQ benchmark across four question types. AutoBioKG shows the strongest overall performance on yes/no, factoid, and list questions, while performance on summary questions remains comparable to the strongest baselines. **b,** Qualitative comparison illustrating the semantic advantages of the composite triple schema. Two case studies demonstrate how AutoBioKG preserves information that is omitted by conventional triples. In Case Study 1 (top), AutoBioKG retains fine-grained entity attributes, including specific mutations such as p.W1095X, that distinguish otherwise similar gene-disease associations. In Case Study 2 (bottom), explicit conditions such as luminal calcium concentration distinguish opposing effects of the same biological interaction.

The advantage of AutoBioKG was most evident on list questions, while it also led on factoid questions. Both tasks depend on retrieving precise details. For factoid questions, AutoBioKG achieved a strict accuracy of 33.0%, 4.0 percentage points above EDC, the strongest baseline. For list questions, the F-measure reached 34.1%, 15.7 percentage points above EDC, the second-best baseline. These results suggest that retaining entity attributes and contextual conditions is useful when questions depend on fine-grained biomedical facts. AutoKG represents knowledge as keyword cooccurrence graphs lacking directed semantic relations and achieved a strict accuracy of 25.0% and a list F-measure of 11.1%. EDC outperformed AutoKG using conventional triples, but this representation does not explicitly retain contextual attributes such as specific gene mutations or dosage information.

Compared with fact-based queries, summary questions require not only relevant evidence but also coherent paragraph-level synthesis. AutoBioKG did not show the same advantage over the baselines on this task, achieving a ROUGE-2 recall of 20.8%, close to the 21.2% obtained by AutoKG, while its ROUGE-2 F1 was 11.6%, slightly below the 12.0% achieved by EDC. This pattern suggests that the gains observed on fact-focused questions are less pronounced for paragraph-level synthesis, where performance also depends on how retrieved evidence is organized and used by the downstream language model.

#### 3.6.2 Case study: Capturing dynamic biomedical associations via context-aware knowledge graph

To qualitatively demonstrate the practical utility of the context-aware knowledge graph, we present two representative case studies based on the BioASQ document corpus, as illustrated in **Figure 4b**. The first case represents a standard benchmark question selected to validate retrieval precision, while the second introduces a complex mechanistic query specifically formulated to assess the system’s ability to resolve context-dependent conflicts.

The first case study involves a factoid query asking which specific mutations of the *SCN5A* gene are implicated in Brugada syndrome. Traditional knowledge graphs, restricted to conventional triples such as (*SCN5A*, related to, *Brugada Syndrome*), often yield non-specific associations that obscure clinically actionable details. In contrast, AutoBioKG populates the graph with specific composite triples, such as (*SCN5A* [attribute: p.W1095X], related to, *Brugada Syndrome*). When processing this query, the system executes precise retrieval operations to enumerate all *SCN5A* mutation attributes associated with the disease, delivering an answer that is not only accurate but also comprehensive and clinically relevant. This example highlights that explicit modeling of entity attributes is critical for resolving granular queries essential for precision medicine.

The second case study involves a mechanistic query regarding the regulation of the Ryanodine Receptor (*RyR*) by *Junctin*, where the interaction is modulated by physiological conditions. When represented as conventional triples, these context-dependent effects appear as conflicting relations, such as (*Junctin*, activates, *RyR*) and (*Junctin*, inhibits, *RyR*). Conversely, AutoBioKG models these interactions as composite triples with explicit conditions: (*Junctin*, activates [condition: low luminal Ca^2+^], *RyR*) and (*Junctin*, inhibits [condition: high luminal Ca^2+^], *RyR*). When querying this relationship, the system disambiguates these opposing effects based on the specific physiological context. This capability enables the knowledge graph to represent context-dependent cellular signaling interactions, thereby extending its utility from fact retrieval to mechanistic inference.

## 4 Discussion

A persistent challenge in biomedical knowledge graph construction is balancing the flexibility of an open relation schema with the contextual specificity needed to represent biological mechanisms. Traditional relation extraction provides structured outputs but is constrained by predefined relation inventories, while open information extraction removes this restriction at the cost of representing most findings as conventional subject-predicate-object triples. AutoBioKG addresses this gap through composite triples. The representation analysis in Fig. 3 makes this information loss explicit. Removing Attribute and Condition fields maps multiple distinct composite triples onto the same conventional triple, whereas retaining these fields preserves their contextual distinctions. Importantly, retaining additional attributes and conditions did not prevent the extractor from generalizing to predefined schemas. The self-trained OIE model improved across all three public relation benchmarks and outperformed the evaluated LLM baselines in zero-shot testing. These results suggest that richer relation structure can be retained without restricting the model to the BioOpenIE schema.

The BioASQ evaluation provides a complementary view of the representation at the graph level. The clearest gain appeared on list questions, while AutoBioKG also outperformed the compared methods on yes/no and factoid questions. Summary questions showed a different pattern. AutoBioKG remained close to the strongest baselines on ROUGE-based summary metrics but did not lead them, suggesting that gains observed on fact-focused questions do not necessarily translate into paragraph-level synthesis. Thus, the downstream results support the value of context-rich graph evidence for factual retrieval, while also showing that answer generation remains an important part of the overall system.

## 5 Conclusion

In this study, we presented AutoBioKG, an end-to-end framework for constructing context-aware biomedical knowledge graphs from scientific literature. By representing entity attributes and relation conditions within composite triples, the framework preserves contextual information that is not explicitly retained in conventional triples. The self-trained OIE model generalized across three public relation schemas and outperformed the evaluated LLM baselines in zero-shot testing. At the graph level, AutoBioKG achieved the strongest overall performance on yes/no, factoid, and list questions in BioASQ, while remaining competitive on summary questions. Together, these results demonstrate the potential of context-aware knowledge representation to improve both biomedical relation extraction and graph-supported question answering. In future work, we aim to extend AutoBioKG beyond standard research articles to document types such as patents and clinical trial reports. We also plan to investigate iterative active learning to improve adaptation to emerging biomedical topics. Ultimately, we envision deploying this framework as a scalable foundation for evidence-based precision medicine and automated scientific discovery.

## Code Availability

The source code for the AutoBioKG framework, including the self-training pipeline and model training scripts, is available at https://github.com/ZYK006/AutoBioKG.

## Acknowledgements

This work was supported by the Major Project of Guangzhou National Laboratory (Grant No. SRPG22-007); the National Key R&D Program of China (Grant No. 2022YFF1202101); the National Key R&D Program of China (Grant No. 2023YFF- 1204701); the National Natural Science Foundation of China (Grant Nos. 12371485 and 82400622); the Major Project of Guangzhou National Laboratory (Grant No. GZNL2025C01013); and the Guangdong Basic and Applied Basic Research Foundation (Grant No. 2025A1515011597).

## A Ablation Analysis of Self-training

To examine the contribution of the main components in self-training, we compared our model with three ablated variants. Our model combines pseudo-labels generated by the multitask seed model with those generated by separately trained single-task models. The ablations remove self-training, the single-task pseudo-label source, or relation completion and verification from the single-task branch. Table 2 reports precision, recall, and F1 on BioOpenIE, DDI, ChemProt, and BioRED.

**Table 2:**
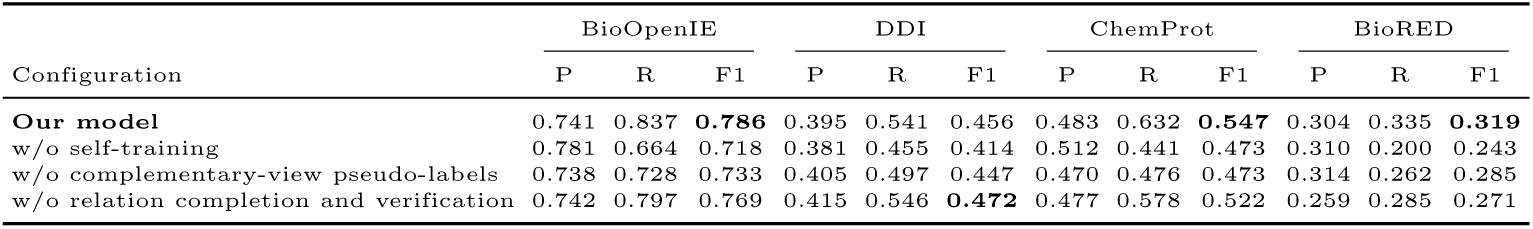
Ablation analysis of self-training. P, R, and F1 denote precision, recall, and F1 score, respectively. “w/o” denotes removal of the indicated component. The highest F1 score for each dataset is shown in bold.

| Configuration | BioOpenIE |  |  | DDI |  |  | ChemProt |  |  | BioRED |  |  |
| --- | --- | --- | --- | --- | --- | --- | --- | --- | --- | --- | --- | --- |
|  | P | R | F1 | P | R | F1 | P | R | F1 | P | R | F1 |
| **Our model** | 0.741 | 0.837 | **0.786** | 0.395 | 0.541 | 0.456 | 0.483 | 0.632 | **0.547** | 0.304 | 0.335 | **0.319** |
| w/o self-training | 0.781 | 0.664 | 0.718 | 0.381 | 0.455 | 0.414 | 0.512 | 0.441 | 0.473 | 0.310 | 0.200 | 0.243 |
| w/o complementary-view pseudo-labels | 0.738 | 0.728 | 0.733 | 0.405 | 0.497 | 0.447 | 0.470 | 0.476 | 0.473 | 0.314 | 0.262 | 0.285 |
| w/o relation completion and verification | 0.742 | 0.797 | 0.769 | 0.415 | 0.546 | **0.472** | 0.477 | 0.578 | 0.522 | 0.259 | 0.285 | 0.271 |

Self-training improved F1 across all four datasets relative to the model without self-training, with gains ranging from 4.2 to 7.6 percentage points. Adding complementary-view pseudo-labels generated by the separately trained single-task models further improved F1 across all four datasets compared with using multitask pseudo-labels alone. The largest additional gains were observed on ChemProt and BioOpenIE, reaching 7.4 and 5.3 percentage points, respectively. These results show that the additional single-task pseudo-label source strengthens self-training across both BioOpenIE and external relation schemas.

Relation completion and verification provided the clearest gains on BioRED and ChemProt, where their inclusion increased F1 by 4.8 and 2.5 percentage points, respectively. BioOpenIE also improved by 1.7 percentage points. Recall increased by 5.0, 5.4, and 4.0 percentage points on BioRED, ChemProt, and BioOpenIE, respectively, indicating broader relation coverage on these datasets. DDI showed a different pattern, with the ablated variant achieving 1.6 percentage points higher F1. These results indicate that the contribution of relation completion and verification varies across relation schemas, with the strongest gains observed on BioRED and ChemProt.

## B Knowledge Graph Retrieval-Augmented Generation Pipeline

Our KG-RAG implementation adapts the framework proposed by Soman et al. [40] with modifications tailored to leverage UMLS-normalized knowledge graph structures. The pipeline consists of four sequential stages: entity extraction, entity matching, context retrieval, and answer generation.

### B.1 Large Language Model Configuration

We employ the August 7, 2025 snapshot of GPT-5 mini [36] for both entity extraction and answer generation through an OpenAI-compatible API endpoint, with a maximum completion-token budget of 4,096.

### B.2 Stage 1: Biomedical Entity Extraction

Given a user query *Q*, we extract relevant biomedical entities using a prompted LLM. The extraction prompt instructs the model to identify entities across multiple biomedical categories including diseases, drugs, genes, proteins, biological processes, anatomical structures, medical procedures, symptoms, and biomarkers.

### B.3 Stage 2: Multi-Strategy Entity Matching

For each extracted entity *e_i_ ∈ E*_extracted_, we employ a cascaded matching strategy to identify corresponding nodes in our UMLS-normalized knowledge graph:

1. **Exact Matching**: String matching against entity names, UMLS preferred names, aliases, and CUI identifiers.
2. **Abbreviation Expansion**: Automatic detection and expansion of abbreviation patterns using regular expression matching.
3. **Semantic Vector Search**: Approximate nearest neighbor search over entity embeddings generated using SapBERT [29], retaining candidates with a vector-store distance *≤* 0.3.

Candidates returned by the matching strategies are deduplicated by CUI and ranked using their corresponding matching scores.

### B.4 Stage 3: Context Retrieval from Knowledge Graph

For each matched entity, we retrieve its local neighborhood from the Neo4j knowledge graph, including entity metadata (preferred name, CUI, semantic types, and description), outgoing and incoming relationships, and relationship attributes such as conditions and provenance documents. The maximum number of relationships retrieved per entity is capped at 80. Retrieved facts are serialized into natural language statements.

To retain question-relevant evidence, each entity-specific context is segmented into sentences and encoded using S-PubMedBert-MS-MARCO [41]. For each entity context *C_j_*, we compute the cosine similarity between the question embedding **v***_Q_* and each context-sentence embedding **v***_si_* :

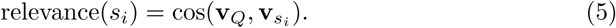

The filtering threshold is determined independently for each entity context as

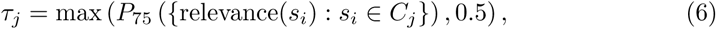

where *P*_75_ denotes the 75th percentile of the similarity scores within that entity context. Sentences with similarity below *τ_j_* are discarded, and the remaining sentences are ranked in descending order of similarity. Given *M* retrieved entity contexts, at most

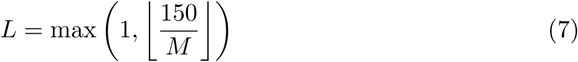

sentences are retained from each entity context. The retained sentences are concatenated to form the final evidence context for answer generation.

### B.5 Stage 4: Answer Generation

The final stage constructs prompts combining retrieved context with question-type-specific instructions. We define four prompt templates corresponding to BioASQ question types (Table **??**).

### B.6 Configuration Parameters

Key hyperparameters used in our KG-RAG pipeline are summarized in Table **??**.

For baseline knowledge graphs (AutoKG, EDC, iText2KG) lacking UMLS normalization, we employ the same pipeline architecture but replace UMLS-specific matching with generic property matching based on available node attributes. AutoKG uses an entity vector-search distance threshold of 0.5, whereas AutoBioKG, EDC, and iText2KG use a threshold of 0.3. The AutoKG threshold was fixed during preliminary pipeline validation before the final BioASQ evaluation because a threshold of 0.3 failed to retrieve usable context for this graph.

### B.7 Computational Infrastructure

All experiments regarding the BioASQ evaluation and KG-RAG inference were performed on the Sugon computing platform. For the inference tasks, the computational resources were allocated with 16 cores of a Hygon 7490 processor and a single BW1000 domestic intelligent acceleration card (32 GB VRAM). This configuration ensured efficient execution of the large language model and dense retrieval operations.

## C Prompt Engineering for Dataset Construction

To construct a high-quality, open-domain relation extraction dataset, we implemented a six-step pipeline driven by Large Language Models (LLMs). This pipeline is designed to support open-world extraction while maintaining biomedical accuracy. Below, we provide the detailed prompts used in each step.

### C.1 Step 1: Semantic Segmentation and Coreference Resolution

The raw introduction text is first segmented into semantic paragraphs to facilitate local context analysis.

##### Prompt for Semantic Segmentation and Coreference Resolution

**Objective:** Segment the following introduction text into 3 to 5 paragraphs. Ensure that each paragraph is concise and focused on a specific topic or research aspect. Preserve the original wording except for coreference replacements, and otherwise only insert paragraph breaks.

**Segmentation Rules:**

1. **Thematic Organization:** Divide the introduction based on major thematic shifts:

- First segment: General background knowledge and introduction of key biological concepts.
- Second segment: Specific mechanisms, processes, and prior research findings.
- Third segment: Research gap and current study’s approach or objectives.
2. **Natural Transitions:** Place segment breaks at natural transition points where the focus shifts.
3. **Pronoun Replacement:** Replace pronouns (e.g., ‘it’, ‘this’, ‘they’) with the corresponding nouns based on context to maintain clarity.
4. **Content Integrity:** Ensure the original text remains unchanged, except for replacing pronouns.

**Output Format:**

[

{"paragraph1": "<TEXT of the first paragraph>"},

{"paragraph2": "<TEXT of the second paragraph>"},

…

]

Text to segment: [Input Text Here]

### C.2 Step 2: Entity Extraction

We extract specific biomedical entities based on a predefined schema to ensure granularity.

##### Prompt for Entity Extraction

Your task is to extract specific biomedical entities from the following scientific text. Adhere to the following requirements:

1. The identified entities should be unique and specific to the biomedical context. Avoid overly general terms (e.g., “gene”, “protein”) unless they refer to specific instances.
2. Only retain the entity name, excluding descriptive modifiers.
3. If both full name and abbreviation appear, include both (e.g., “Tumor Necrosis Factor (TNF)”).
4. **Allowed Entity Types:** [“Clinical Drug”, “Disease or Syndrome”, “Gene or Genome”, “Protein”, “Biologic Function”, “Cell”, “Cell Component”, “Tissue”, “Enzyme”, “Hormone”, “Vitamin”, “Receptor”, “Virus”, “Lipid”, . . .].

**Output Format:**

[

{

"entity_name": "<THE name of the entity>",

"entity_type": "<THE type of the entity>",

"entity_description": "<BRIEF description>"

}

]

### C.3 Step 3: Composite Triple Extraction

Using the entities identified in Step 2, we extract relationships within each paragraph.

##### Prompt for Composite Triple Extraction

You have identified the following biomedical entities from the previous text. Your task is to extract **only meaningful biomedical relationships** between these entities. **List of Entities:** [List of Entities from Step 2]

**Relationship Extraction Criteria:**

1. **Entity and Relationship Representation:**

- **Entity Attributes (Entity:Attribute):** Apply only when the attribute is scientifically valuable (e.g., “BRCA1:c.5266dupC”). Discard general attributes.
- **Relationship Conditions (Relation:Condition):** Apply when the text specifies contextual factors (e.g., “increases activity of:under high temperature”).
2. **Validation Criteria:**

- Valid: Must reflect biologically significant interactions (e.g., “activates”, “biomarker”).
- Invalid: Exclude attribute relationships (“is a”), experimental setups, or uncertain statements (“may cause”).
- Handling Indirect Relationships: Use “indirect-xx” format if the intermediate step is not explicitly mentioned.

**Output Format:**

[

{

"head_entity": "Head Entity" or "Head Entity:Attribute",

"relation_type": "Relation" or "Relation:Condition",

"tail_entity": "Tail Entity" or "Tail Entity:Attribute"

}

]

### C.4 Step 4: Paragraph-level Triple Verification

Extracted triples are immediately verified to reduce hallucinations and ensure format compliance.

##### Prompt for Paragraph-level Triple Verification

**Task:** Validate each triple and decide to **keep**, **correct**, or **discard**.

**Processing Rules:**

1. **Judgment type:**

- Keep: triple is valid.
- Correct: Fix errors in entities, states, or relation types based on text.
- Discard: Invalid or hallucinatory triple.
2. **Validation Criteria:**

- Entities: Verify attributes add scientific value. Base entities must come from the provided list.
- Relationships: Must reflect biologically significant interactions. Concepts like ‘risk’ or ‘level’ should be part of the relationship (e.g., “increases risk of”), not entity attributes.

**Output Format:**

[

{

"Triple": "(entity, relation, entity)",

"decision": "keep/discard/correct",

"corrected result": "…",

"reason_for_triple": "Explain the basis for judgment"

}

]

### C.5 Step 5: Cross-paragraph Triple Completion

After paragraph-level extraction, we scan the text for relationships that span across sentences.

##### Prompt for Cross-paragraph Triple Completion

Identify cross-paragraph triples overlooked during initial extraction. Focus only on relationships where entities appear in different sentences.

**Guidelines:**

1. Combine information across sentences using co-references (pronouns, synonyms) or logical connections.
2. Exclude triples already captured in the “Extracted triples” section.
3. Exclude attribute relationships and unsupported assumptions.

**Input:** List of Entities, Existing triples, and Original Text.

**Output Format:** If no additional triples are found, output: “No additional triples beyond the extracted ones were found.” Otherwise, provide the triples in strict JSON format.

### C.6 Step 6: Document-level Triple Verification

Finally, we merge the triples from Steps 4 and 5 and perform a final consistency check against the full Introduction.

##### Prompt for Document-level Triple Verification

**Task:** Using the “List of Entities” and extracted triples, validate each triple and decide to **keep**, **correct**, or **discard**.

*(Note: The detailed criteria for entity attributes, composite relationships, and correction guidelines are identical to Step 4 to ensure consistency across the pipeline.)*

**Correction Guidelines:**

- **Entity Normalization:** Expand abbreviations to “Full Name (Abbreviation)” format.
- Corrections must use direct text evidence without external information.
- Ensure corrected triples remain consistent with the original text.

**Text to analyze:** [Full Introduction Text]

**Table 5:** Entity Type Definitions. This table provides the definitions for the 23 core biomedical entity types used for automated annotation and information extraction in this study. The schema is derived and adapted from the UMLS Semantic Network to ensure broad coverage of key concepts in biomedical literature.

| # | Entity Type | Definition |
| --- | --- | --- |
| 1 | Clinical Drug | A substance approved for use in the diagnosis, treatment, or prevention of disease in humans or animals. |
| 2 | Disease or Syndrome | A pathological condition characterized by a specific set of signs and symptoms, representing a disorder of structure or function. |
| 3 | Gene or Genome | A distinct sequence of nucleotides in DNA or RNA, which can be a functional unit for inheritance. This category also includes the entire genetic material of an organism. |
| 4 | Protein | A large biomolecule composed of one or more long chains of amino acid residues, performing a vast array of functions within organisms. |
| 5 | Biologic Function | A process or set of processes carried out by a cell or organism. This specifically includes metabolic or signaling pathways, which are series of chemical or molecular interactions that govern cellular responses. |
| 6 | Cell | The fundamental structural and functional unit of all known living organisms. |
| 7 | Cell Component | A constituent part of a cell, such as an organelle (e.g., nucleus, mitochondrion) or a macromolecular complex. |
| 8 | Tissue | An ensemble of similar cells and their extracellular matrix from the same origin that together carry out a specific function. |
| 9 | Biomedically Active Substance | A general category for any chemical that has a demonstrable biological effect on an organism, including endogenous metabolites and exogenous compounds. |
| 10 | Enzyme | A protein that acts as a biological catalyst to accelerate chemical reactions. |
| 11 | Hormone | A signaling molecule produced by glands in multicellular organisms that is transported by the circulatory system to target distant organs to regulate physiology and behavior. |
| 12 | Vitamin | An organic compound and a vital nutrient that an organism requires in limited amounts for its metabolism. |

**Table 5 – continued from previous page**
| # | Entity Type | Definition |
| --- | --- | --- |
| 13 | Prostaglandin | A group of physiologically active lipid compounds having diverse hormone-like effects in animals. |
| 14 | Immunologic Factor | A substance that is part of the immune system and plays a role in the immune response, such as a cytokine, antibody, or complement component. |
| 15 | Pharmacologic Substance | A chemical substance used in the treatment, cure, prevention, or diagnosis of disease or used to otherwise enhance physical or mental well-being. It is a broader category than Clinical Drug, including experimental compounds. |
| 16 | Receptor | A protein molecule, embedded in either the plasma membrane or the cytoplasm of a cell, to which one or more specific kinds of signaling molecules may attach. |
| 17 | Cell or Molecular Dysfunction | An abnormal or impaired function at the cellular or molecular level, often leading to or characteristic of a disease state. |
| 18 | Cell Function | The normal physiological activities performed by a cell, such as cell growth, differentiation, apoptosis, or signal transduction. |
| 19 | Molecular Function | The elemental activities of a gene product at the molecular level, such as binding to a molecule or catalysis of a biochemical reaction. |
| 20 | Virus | A submicroscopic infectious agent that replicates only inside the living cells of other organisms. |
| 21 | Neuroreactive Substance or Biogenic Amine | A chemical substance that has a specific effect on the nervous system, including neurotransmitters (e.g., dopamine, serotonin) and neuromodulators. |
| 22 | Steroid | A biologically active organic compound with four rings arranged in a specific molecular configuration, serving as hormones, vitamins, or structural components of cell membranes. |
| 23 | Lipid | A class of organic molecules including fats, waxes, and sterols, which are integral to cell structure, energy storage, and signaling. |

